# IGF1 peptide targets Rett Syndrome astrocytes to degrade IGF binding protein, rescue synaptogenesis and restore mitochondrial function

**DOI:** 10.64898/2026.01.22.701190

**Authors:** Prachi Ojha, Velina Kozareva, Alexandria Barlowe, Fabian Schulte, Ruby Minh Lam, Danielle L Tomasello, Ernest Fraenkel, Rudolf Jaenisch, Mriganka Sur

## Abstract

Rett syndrome (RTT), a severe neurodevelopmental disorder caused by mutations in MECP2, leads to profound synaptic and circuit deficits in the brain. While neurons have historically been the focus of RTT pathology, emerging evidence implicates astrocytes in non-cell autonomous mechanisms that impair synaptic structure, function and development. Here, we uncover a central role for astrocyte-secreted IGFBP2 in mediating these deficits and demonstrate that treatment with an IGF1-derived peptide restores synapse formation by promoting IGFBP2 degradation. Using an indirect astrocyte-neuron co-culture system, we show that astrocytes derived from RTT model mice suppress excitatory synapse formation in wild-type neurons and that this impairment is reversed when RTT astrocytes are treated with IGF1(1-3) peptide. Proteomic analysis reveals elevated levels of IGFBP2 in RTT astrocytes and their conditioned media. IGF1(1-3) peptide treatment leads to proteasomal degradation of IGFBP2, increasing IGF1 bioavailability, restoring mitochondrial function, and enhancing downstream PI3K/Akt signaling in neurons. Our data define a molecular mechanism by which astrocyte dysfunction in RTT can be rescued and provide a mechanistic basis for the therapeutic efficacy of IGF1(1-3) peptide, including Trofinetide, an FDA-approved IGF1 peptide mimetic, in RTT.

**Significance Statement:** Astrocyte dysfunction is increasingly recognized as a contributor to neurodevelopmental disorders, yet their precise mechanisms of action remain elusive. Here, we identify IGFBP2 as a key astrocyte-derived inhibitor of synaptogenesis in Rett syndrome. We show that an IGF1-derived peptide, IGF1(1-3), depletes IGFBP2 via proteasomal degradation. This restores IGF1 bioavailability and rescues synaptic function in a non-cell-autonomous manner. These findings provide a mechanistic explanation for the clinical efficacy of IGF1 peptide and its mimetics in Rett syndrome, and highlight astrocytes as rational therapeutic targets in neurodevelopmental and other disorders.

## Introduction

Rett syndrome (RTT) is a severe neurodevelopmental disorder caused by mutations in the X-linked gene *MECP2* (1, 2), encoding methyl-CpG binding protein, which binds to DNA widely across the genome. Originally described as a transcriptional repressor (3, 4), MeCP2 is now recognized to have diverse functions (5, 6), including roles as transcriptional activator as well as repressor (7–9). The loss of MeCP2 thus has pleiotropic effects, making a functional analysis of its downstream pathways essential for devising therapeutic strategies. A pivotal discovery was that RTT is a reversible disorder: restoring MeCP2 expression even after symptom onset can rescue neurological function in mouse models, indicating that underlying synaptic circuits remain immature but plastic and potentially amenable to maturation (10, 11). In studies of activity-dependent cortical plasticity, we identified insulin-like growth factor 1 (IGF1) signaling as a key pathway that promotes synaptic maturation during development: reduced IGF1 signaling accompanies visual deprivation-induced plasticity of neuronal responses and circuits in visual cortex, whereas systemic application of IGF1(1-3), an active tripeptide fragment of IGF1, upregulates IGF1 signaling, promotes synaptic maturation, and prevents such plasticity (12). We subsequently showed that systemic administration of IGF1(1-3), or of full-length IGF1, improves synaptic maturation, spine density, circuit function, and behavior in Mecp2 null mice (13–17). These findings provided the foundation for clinical trials of IGF1 and IGF1 peptide in RTT, including trials with Trofinetide - a mimetic of IGF1(1-3) with an added methyl group making it resistant to breakdown and prolonging its bioavailability - leading to the recent FDA approval of Trofinetide as the first disease-modifying therapeutic for RTT (18, 19). However, the cellular and molecular mechanisms by which IGF1(1-3) peptide or Trofinetide exert their therapeutic effects remain incompletely understood (20).

While RTT was initially considered a neuron-intrinsic disorder, astrocytes are now recognized as critical contributors to disease pathophysiology (21–23). Conditioned medium from MeCP2-deficient astrocytes fails to support synapse formation in wild-type neurons, indicating the presence of astrocyte-derived factors that actively impair synaptic connections and neuronal function through non-cell autonomous mechanisms (24–28). Furthermore, astrocyte conditioned media from mouse models of RTT show upregulation of IGF binding proteins (29), suggesting a mechanism for decreased IGF1 signaling in RTT. Thus, MeCP2 deficiency alters the expression of a range of astrocyte-secreted molecules that influence neurons, and reversing these changes presents a compelling potential target for therapeutic strategies for RTT.

Here, we identify insulin-like growth factor binding protein 2 (IGFBP2) as a major astrocyte-secreted inhibitor of synaptogenesis in RTT. We show that IGFBP2 is upregulated in RTT astrocytes, and that treatment with IGF1(1-3) peptide induces its proteasome-dependent degradation. IGFBP2 binds to and sequesters IGF1, and its degradation restores IGF1 bioavailability, upregulates PI3K/Akt signaling in neurons, and rescues synaptic protein expression. Furthermore, our proteomic profiling reveals a previously unrecognized connection to mitochondrial dysfunction: while IGF1 primarily interacts with mitochondrial proteins in healthy astrocytes, its interactome in RTT astrocytes shifts to proteasomal components. IGF1(1-3) peptide treatment reverses the associated mitochondrial deficits, restoring metabolic support for neurons. Our findings define an astrocyte-centered mechanism for the effectiveness of IGF1(1-3) peptide and Trofinetide, linking IGFBP2 degradation to the restoration of both synaptic signaling and mitochondrial function, and identify this mechanism as a promising therapeutic node for RTT and other neurodevelopmental disorders.

## Results

### MeCP2-deficient astrocytes impair excitatory synapse formation via non-cell-autonomous mechanisms

To investigate the contribution of astrocytes to RTT pathogenesis, we established a neuron-astrocyte co-culture system using combinations of primary mouse wild-type and Mecp2^tm1.1Bird^/J knockout astrocytes (WT-A, KO-A) and neurons (WT-N, KO-N) (Fig. 1a). This system allowed us to independently manipulate astrocyte and neuronal genotype and thereby isolate the effects of astrocytes on neuronal synapse formation.

**Fig. 1.**
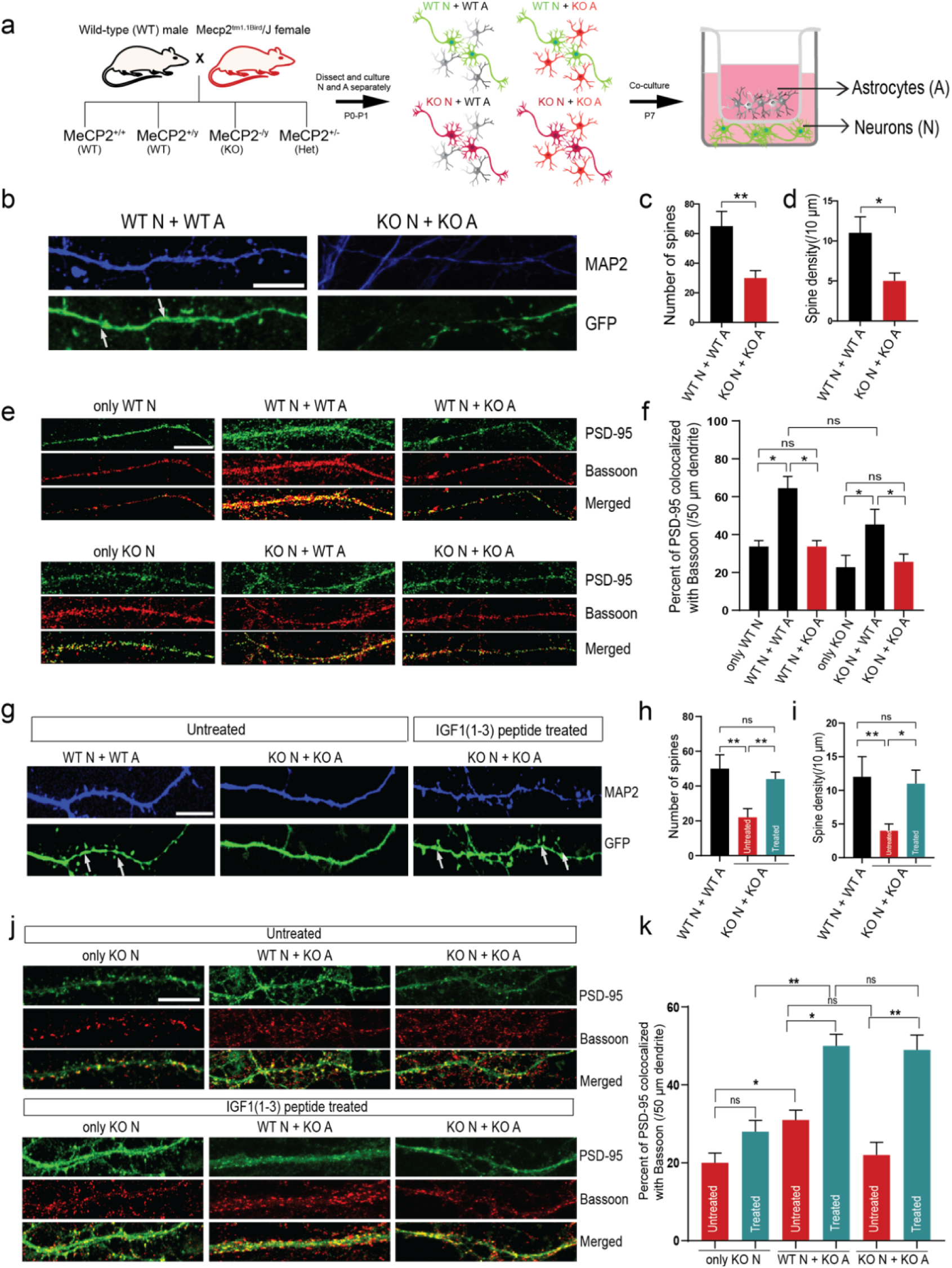
MeCP2-deficient astrocytes impair excitatory synapse formation in neurons, which is rescued by IGF1(1-3) peptide treatment. (a) Schematic of the neuron-astrocyte co-culture system using wild-type (WT) or MeCP2 knockout (KO) neurons (N) and astrocytes (A), and experimental timeline. (b) Representative images of 50 μm segments of dendrites from WT N + WT A and KO N + KO A co-cultures stained for MAP2 (blue) and GFP (green) (scale bar = 10 μm). White arrows mark representative dendritic spines. (c-d) Quantification of total spine number (c) and spine density (spines/10 μm dendrite) (d). KO N + KO A co-cultures show reduced spine numbers and density compared to WT N + WT A (**p < 0.01, *p < 0.05; one-way ANOVA with Tukey’s post hoc test; n = 4 cultures). (e) Representative images showing Bassoon (presynaptic marker, red), PSD-95 (postsynaptic marker, green), and merged channels in dendrites (scale bar = 10 μm). (f) Quantification of PSD-95/Bassoon colocalization (% per 50 μm dendrite; *p < 0.05; ns, non-significant; one-way ANOVA with Tukey’s post hoc test; n = 6 cultures). (g) Representative images of MAP2 (blue) and GFP (green) in WT N + WT A and KO N + KO A, where KO A are untreated or treated with IGF1(1-3) peptide (100 ng/mL, 24 h). (h-i) Quantification showing that peptide treatment rescues spine number (h) and spine density (i) in KO N + KO A (**p < 0.01, *p < 0.05 vs. untreated KO; one-way ANOVA with Tukey’s post hoc test). (j) Representative dendrite images from WT N + KO A and KO N + KO A co-cultures, untreated and peptide-treated (scale bar = 10 μm). (k) Quantification of excitatory synapse number in KO A co-cultures with or without peptide treatment (*p < 0.05, **p < 0.01 vs. untreated KO; one-way ANOVA with Tukey’s post hoc test). All co-culture combinations were tested; representative conditions are shown for clarity.

We first examined dendritic spine structure in matched genotype co-cultures. Neurons from KO mice cultured with KO astrocytes exhibited a significant reduction in dendritic spine number and density compared to WT-N + WT-A cultures (Fig. 1b-d), indicating that Rett syndrome model co-cultures display structural synaptic abnormalities. However, because both neurons and astrocytes differ in these conditions, this experiment does not distinguish whether these defects arise from neuronal, astrocytic, or combined contributions.

To characterize astrocytic MeCP2 status in this system, we measured MeCP2 protein expression in cultured astrocytes. WT astrocytes expressed higher levels of MeCP2 compared to heterozygous astrocytes, whereas KO astrocytes lacked detectable MeCP2 expression (SI Appendix, Fig. S1a, b). Astrocyte morphology was also altered in KO-A + KO-N co-cultures (SI Appendix, Fig. S1c, d), consistent with cellular changes accompanying MeCP2 loss.

We next asked whether astrocytes specifically contribute to the observed synaptic abnormalities. To this end, we quantified the colocalization of the excitatory postsynaptic marker PSD-95 with the presynaptic marker Bassoon across all neuron-astrocyte combinations. WT neurons cultured with KO astrocytes exhibited a ∼50% reduction in PSD-95/Bassoon colocalization compared to WT-N cultured with WT-A controls (33.7 ± 3.2% vs. 64.5 ± 6.2%) (Fig. 1e, f), demonstrating that KO astrocytes are sufficient to impair excitatory synaptogenesis in otherwise wild-type neurons. This impairment was further exacerbated in KO-N + KO-A co-cultures (25.6 ± 4.1%) and was partially rescued when KO neurons were cultured with WT astrocytes (45.4 ± 7.9%) (Fig. 1e, f), indicating that both astrocytic and neuronal MeCP2 contribute to synaptic deficits, with astrocytes exerting a strong non-cell-autonomous effect. Consistent with these findings, we found a significant reduction in total PSD-95 levels in KO neurons (SI Appendix, Fig. S1g, h). In contrast to excitatory synapses, inhibitory synapse formation, assessed by Gephyrin/Bassoon colocalization, was unchanged across all co-culture conditions (SI Appendix, Fig. S1e, f), indicating that MeCP2 deficiency selectively affects excitatory synaptogenesis.

Together, these results indicate that structural synaptic abnormalities observed in Rett syndrome are accompanied by a selective impairment in excitatory synapse formation driven importantly by MeCP2-deficient astrocytes.

### IGF1(1-3) peptide treatment of astrocytes restores synapse numbers in neurons

To test whether IGF1(1-3) peptide can rescue synaptic impairments caused by KO astrocytes, we treated astrocytes with the peptide prior to co-culture with neurons (see Methods). Because our previous study (13) only tested the efficacy of this peptide *in vivo* and not in cultures, we first performed an *in vitro* dose-response assay to determine the appropriate concentration of the peptide that activates broad downstream signaling pathways such as phospho-ERK (p-ERK). All three doses of the peptide (50 ng/mL, 100 ng/mL, and 200 ng/mL) were effective in increasing p-ERK levels in KO astrocytes (SI Appendix, Fig. S1i, j). We thus selected the intermediate concentration (100 ng/mL) for subsequent experiments.

We found that WT neurons co-cultured with KO astrocytes treated with IGF1 peptide exhibited significantly increased dendritic spine density and synapse numbers compared to untreated KO astrocyte co-cultures (Fig. 1g-i). Excitatory synapse formation (PSD-95/Bassoon colocalization) was also significantly restored in neurons exposed to peptide-treated KO astrocytes, reaching levels comparable to WT astrocyte co-culture conditions (55 ± 3.2% for WT-N + KO-A treated vs. untreated, 50.2 ± 4.1% for KO-N + KO-A treated vs. untreated) (Fig. 1j, k). Thus, peptide treatment of KO astrocytes is sufficient to restore their ability to support excitatory synaptogenesis and neuronal spine formation.

Consistent with the previously described lack of effect of KO-As on inhibitory synapse formation, peptide treatment of astrocytes had no effect on inhibitory synapses (SI Appendix, Fig. S1k, l). These data demonstrate that IGF1(1-3) peptide specifically restores excitatory synapse formation and neuronal spine formation by acting on KO astrocytes. To control for the possibility that IGF1 peptide in the media could diffuse from the insert membrane and act directly upon neurons, we also treated WT-N and KO-N with IGF1 peptide and found no change in the number of excitatory or inhibitory synapses, indicating that the observed rescue requires astrocytes (Fig. 1j, k; SI Appendix, Fig. S1m). Thus, IGF1 peptide acts on astrocytes, but not directly on neurons, to rescue excitatory synapse formation in neurons.

### IGFBP2 is upregulated in KO astrocytes and their secretome

To identify secreted astrocytic factors responsible for reduced numbers of excitatory synapses in KO neurons, we performed quantitative proteomic profiling of both astrocyte-conditioned media (ACM) and whole-cell lysates from WT and KO astrocytes (WT-A and KO-A, Fig. 2a). This approach allowed us to examine both extracellular and intracellular alterations associated with MeCP2 loss.

**Fig. 2.**
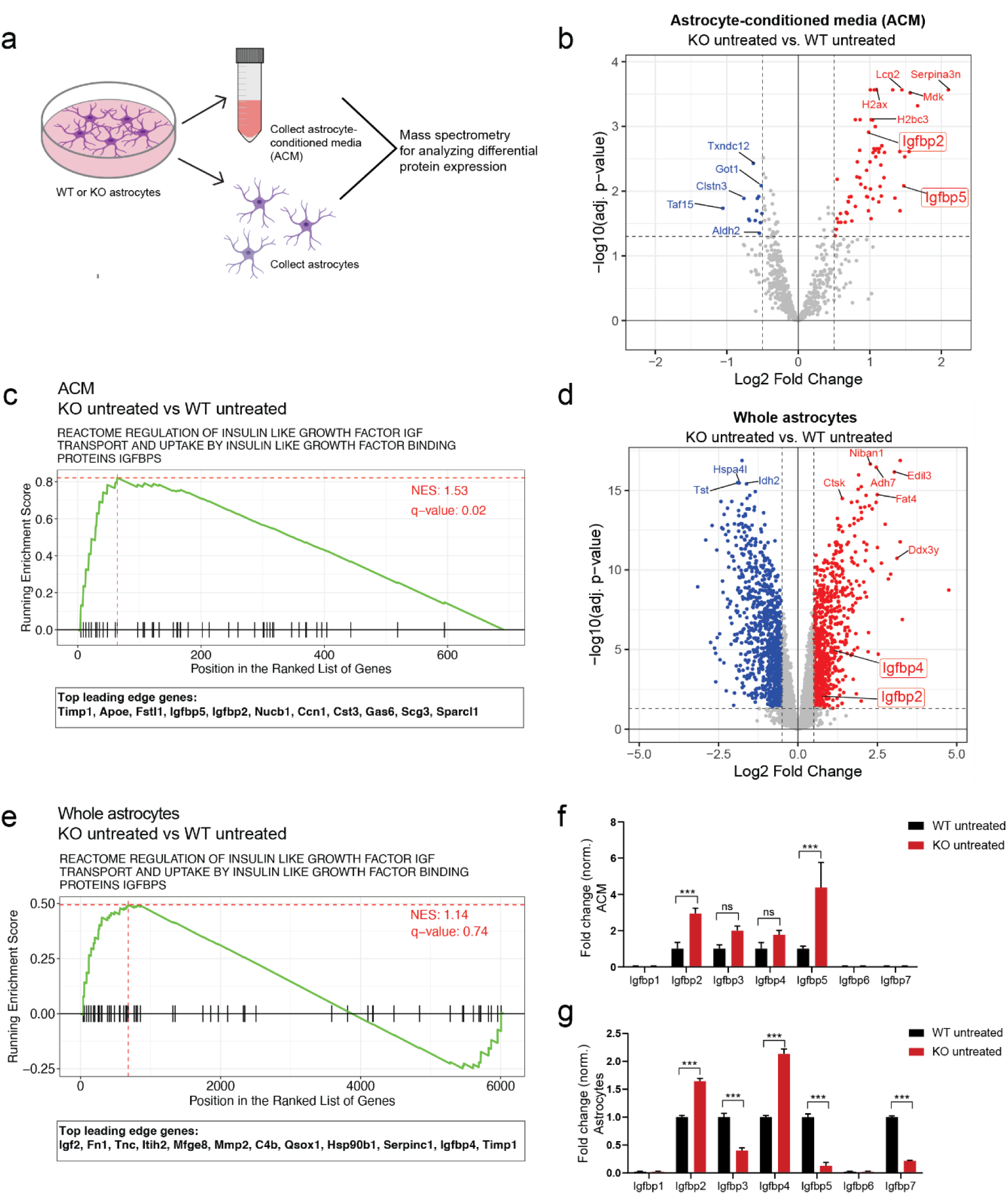
IGFBPs are dysregulated in both MeCP2 KO astrocytes and astrocyte-conditioned media (ACM). (a) Workflow for mass spectrometry analysis of astrocyte-conditioned media (ACM) and cell lysates. (b) Volcano plot of differentially expressed proteins (DEPs) in ACM (KO vs. WT), highlighting upregulated proteins in red and downregulated proteins in blue. Note that Igfbp2 and Igfbp5 highlighted in the red box are upregulated (log₂FC > 0.5, adj. p < 0.05). (c) Gene set enrichment analysis shows significant positive enrichment of IGFBP-related pathway in KO ACM (NES = 1.53, q-val = 0.02). Leading edge genes driving this enrichment are indicated below. Dotted red lines indicate maximal magnitude enrichment score computed across proteins ranked by degree of differential expression in indicated comparison. Positive ES score indicates relative up-regulation of proteins in indicated pathway, corresponding to overrepresentation of pathway proteins (black ticks) at top of ranked protein list. (d) Volcano plot of DEPs in astrocytes (KO vs. WT), highlighting upregulated IGFBPs (red box; log₂FC > 0.5, adj. p < 0.01). (e) GSEA shows non-significant but still positive enrichment of IGFBP-related pathway in KO astrocytes (NES = 1.14, q-val = 0.74). Leading edge genes driving this enrichment are indicated below. (f, g) Fold change of IGFBPs in ACM (f) and astrocytes (g) (***, p < 0.001 vs. WT; ns, non-significant; two-tailed t-test). NES = normalized enrichment score.

Several insulin-like growth factor binding proteins (IGFBPs) were upregulated in both the media and cell lysates of KO astrocytes. In ACM, IGFBP2 and IGFBP5 were among the most significantly increased proteins (log_2_FC = 0.98, adj. p = 1.2 × 10^-3^; log_2_FC = 1.47, adj. p = 8.3 × 10^-3^, respectively) (Fig. 2b, f and SI Appendix, Dataset S1). IGFBP2 was also elevated in whole-cell lysates (log_2_FC = 0.74, adj. p = 7.8 × 10^-3^). In addition, IGFBP4 was upregulated in KO astrocytes (log_2_FC = 1.11, adj. p = 1.02 × 10^-5^) (Fig. 2d, g and SI Appendix, Dataset S2). Together, these changes indicate a shared dysregulation of IGFBP levels and potentially of IGF1 signaling.

To determine the major pathways affected, we performed gene set enrichment analysis (GSEA) of ACM and astrocytic proteomes. GSEA revealed significant positive enrichment of an IGFBP-dependent pathway (Reactome) in ACM and a similar, though not statistically significant, enrichment in astrocytes (Fig. 2c, e and SI Appendix, Fig, S2a, b; Datasets S3, S4).

Beyond IGF signaling, GSEA of KO ACM also showed activation of extracellular matrix (ECM)-related pathways (normalized enrichment score, NES = 1.51, FDR q = 0.002) and suppression of RNA metabolism (Reactome) (NES = −2.11, FDR q = 0.0009) (SI Appendix, Fig. S2a and Dataset S3). Likewise, GSEA of whole-cell astrocytic proteomes revealed dysregulation of metabolic processes, including mitochondrial matrix and cellular respiration, suggesting that MeCP2 deficiency also affects astrocyte metabolism (SI Appendix, Fig. S2b and Dataset S4). Notably, GSEA of secreted proteins also indicated suppression of nervous system development and Robo signaling pathways, which could impact neuronal synapse formation and maturation (SI Appendix, Fig. S2a).

To validate the proteomics results, we performed qPCR for multiple IGFBPs and synaptogenic genes. Igfbp1, Igfbp4, Igfbp5, Igfbp6, as well as EphA3 and Tnf, were significantly upregulated in KO astrocytes, whereas Igfbp2 mRNA was not elevated (SI Appendix, Fig. S2c). This discrepancy between mRNA and protein levels suggests that IGFBP2 is regulated post-translationally and that MeCP2 loss leads to an accumulation of the IGFBP2 protein. Collectively, these results identify IGFBP2 as a key protein upregulated in both ACM and astrocytes.

### Peptide treatment reverses dysregulated proteins in MECP2-deficient astrocytes

To determine whether IGF1 peptide could reverse astrocytic proteome dysregulation, we performed differential proteomic analyses of KO astrocytes and ACM after 24 h of treatment with IGF1(1-3) peptide (100 ng/mL) (Fig. 3a). Comparison of protein fold changes between peptide-treated KO vs. untreated KO astrocytes, and untreated KO vs. WT astrocytes, revealed that most significant proteomic alterations in KO astrocytes were at least partially reversed (75%, 2406/3216 significant differentially expressed proteins, DEPs, from KO vs. WT in the green quadrants), with a strong negative correlation among all DEPs (R = −0.57, p < 0.001) (Fig. 3b). In contrast, ACM showed a smaller fraction of reversed proteins (52%, 359/691 total detected proteins in the green quadrants) and a weaker overall correlation (R = −0.08, p < 0.01) (Fig. 3c), indicating that the peptide primarily acts on intracellular rather than secreted protein networks (SI Appendix, Datasets S1 and S2).

**Fig. 3.**
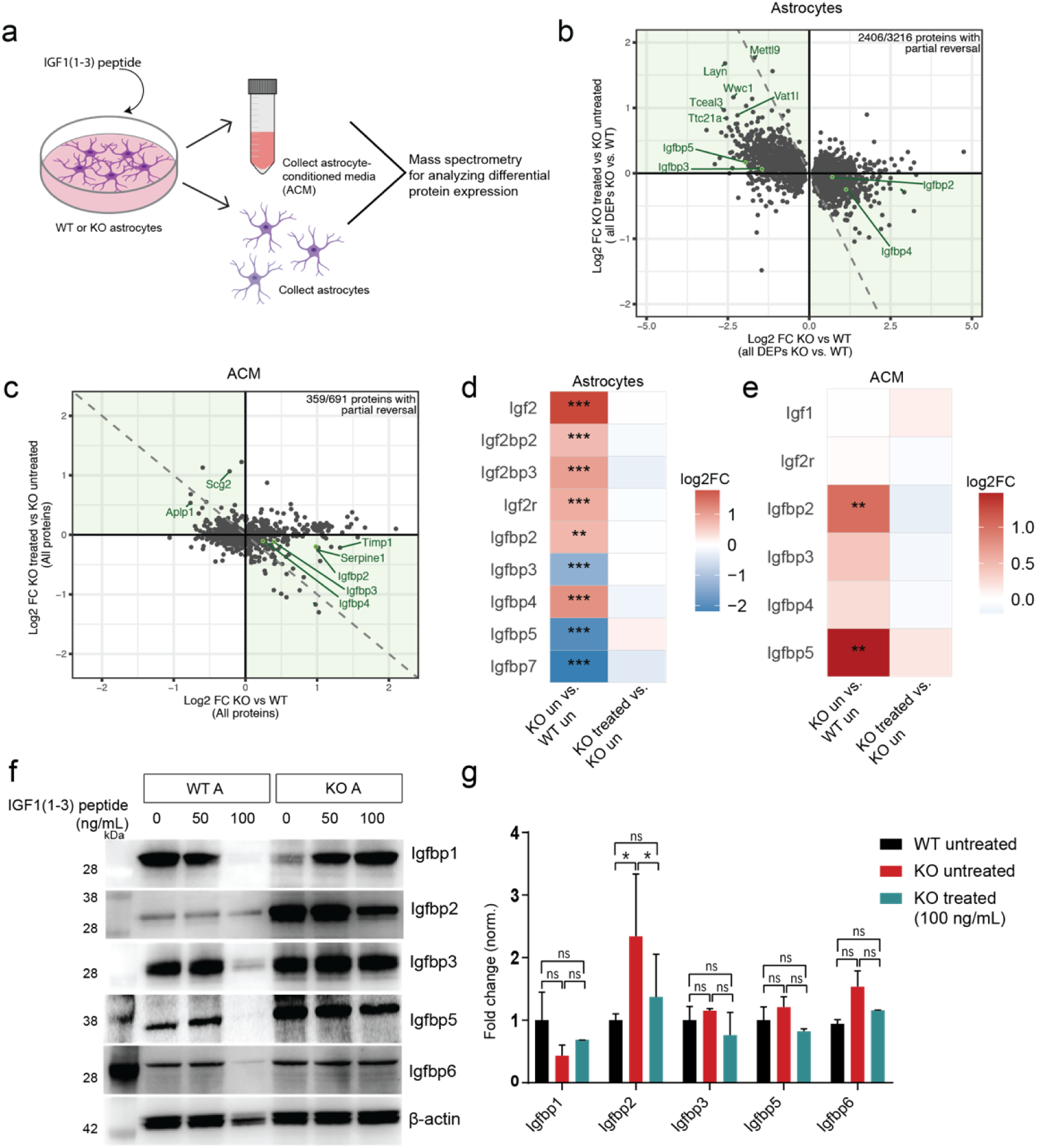
IGF1(1-3) peptide treatment partially reverses dysregulated proteins in MeCP2-deficient astrocytes. (a) Experimental design for peptide treatment and proteomics. (b) Scatter plot of log₂FC (KO + peptide/KO) vs. log₂FC (KO/WT) in astrocytes for all DEPs in KO vs. WT (n = 3216), showing partial reversal for 2406/3216 proteins and strong anti-correlation (Pearson r = −0.57, p < 0.001). Reversed proteins fall in the green quadrants. Top left quadrant shows proteins that are decreased in KO untreated but increased in KO treated and the bottom right quadrant shows proteins that are increased in KO untreated but decreased in KO treated). (c) Scatter plot of log₂FC (KO + peptide/KO) vs. log₂FC (KO/WT) in ACM for all detected proteins (n = 691), showing partial reversal for 359/691 proteins (Pearson r = −0.08, p < 0.01). (d, e) Heatmaps of expression changes in IGF family proteins in astrocytes (d) and in ACM (e); proteins with statistically significant changes are indicated (log₂FC, KO vs. WT and KO+peptide vs. KO), * FDR < 0.05; ** FDR < 0.01; *** FDR < 0.001. (f) Western blots show downregulation of upregulated IGFBP2 in treated vs. untreated KO astrocytes. (g) Quantification of Western blots following treatment with 100 ng/mL peptide (n = 3; *, p < 0.05; two-tailed t-test). Lane 1 or WT untreated was used as a control for all comparisons.

Because IGFBPs emerged as a shared class of upregulated proteins in both astrocytes and ACM, suggesting a coordinated alteration of IGF signaling, we next examined how IGF1(1-3) peptide treatment affected IGF pathway proteins. To visualize these changes, we extracted all detected IGF1 family members from the proteomics dataset and plotted their expression change profiles upon peptide treatment in astrocytes and ACM. In KO astrocytes, many more IGF pathway proteins, including IGFBP2 and IGFBP4, showed reversal after peptide treatment compared to ACM (Fig. 3d, e).

The impact of IGF1(1-3) peptide extended beyond individual IGFBPs. GSEA of treated vs. untreated ACM revealed partial restoration of several synaptic and IGF1 signaling pathways, although none reached statistical significance (SI Appendix, Dataset S5). In contrast, pathway reversal in KO astrocytes was more extensive and included several processes that did reach significance (SI Appendix, Dataset S6).

To summarize these trends, we selected our pathways of interest and plotted them side by side for each condition based on their absolute NES sum (SI Appendix, Fig. S3a-c and Dataset S5, S6). Consistent with the proteomic reversal plots, astrocytes displayed a broader restoration of neuron-synaptogenic and insulin/IGF-related pathways (SI Appendix, Fig. S3b, c), whereas ACM showed only limited recovery (SI Appendix, Fig. S3a). Note that in astrocytes, the IGFBP-related pathway that was earlier positively enriched was now reversed (SI Appendix, Fig. S3b).

Complementary western blot analyses reinforced these findings: IGFBP2 levels were significantly upregulated in KO astrocytes and were strongly reduced following IGF1(1-3) peptide exposure, while other IGFBPs showed minimal changes (Fig. 3f, g). Importantly, IGFBP levels in KO neurons remained unchanged before and after peptide exposure (SI Appendix, Fig. S3d, e), indicating that IGF1(1-3) peptide selectively targets astrocytic rather than neuronal IGFBP dysregulation in RTT.

Collectively, these results demonstrate that MeCP2-deficient astrocytes show aberrant upregulation of IGFBP2 in both intracellular and secreted compartments, and that IGF1(1-3) peptide treatment partially restores the astrocytic proteome by reversing IGFBP2 accumulation and rebalancing key IGF- and synapse-related pathways.

### IGF1(1-3) peptide promotes proteasomal degradation of IGFBP2 and increases IGF1 bioavailability

IGFBP2 is known to bind IGF1 with high affinity, restricting its availability for receptor activation and thereby modulating downstream PI3K-Akt signaling (30–32). To investigate the mechanism underlying IGFBP2 regulation in RTT astrocytes, we tested whether IGF1(1-3) peptide affects the IGFBP2-IGF1 interaction. Co-immunoprecipitation (Co-IP) using FLAG-tagged full-length IGF1 (Fig. 4a) revealed that IGFBP2 binding to IGF1 was markedly elevated in KO astrocytes compared to WT (3.9 ± 0.5-fold over WT), and this binding was significantly reduced following peptide treatment (1.8 ± 0.7) (Fig. 4b, c). These findings indicate that IGF1(1-3) partially competes with IGFBP2 for IGF1 binding, thereby displacing the complex and potentially increasing IGF1 bioavailability for receptor engagement.

**Fig. 4.**
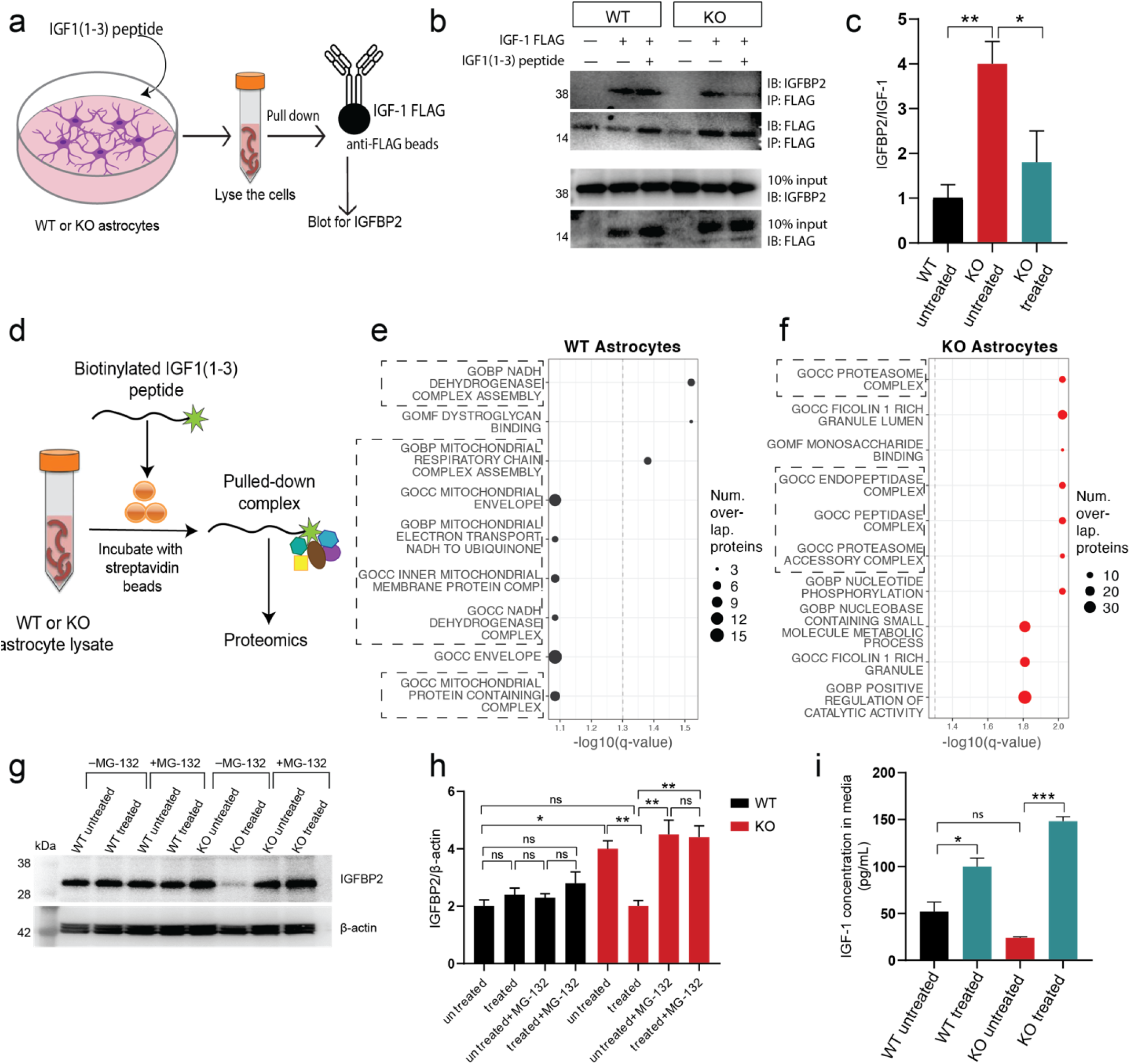
Mechanism of IGF1 peptide action involves IGF1-IGFBP2 displacement and proteasomal degradation of IGFBP2, increasing IGF1 bioavailability. (a) Co-immunoprecipitation experimental schematic using FLAG-tagged IGF1. (b) Representative Western blot of Co-IP samples probed for IGFBP2. (c) Quantification of IGFBP2 binding to IGF1 (n=3; **, p < 0.01; *, p < 0.05). (d) Workflow for biotinylated IGF1(1-3) peptide pull-down and mass spectrometry. (e, f) Gene Ontology (GO) term analysis of proteins enriched in biotinylated IGF1(1-3) pull-downs from WT astrocytes (mitochondrial, respiration terms indicated) (e) and KO astrocytes (proteasome, peptidase terms indicated) (f). (g) Western blot of IGFBP2 levels in KO astrocytes treated with IGF1(1-3) peptide in the presence or absence of proteasome inhibitor MG-132. (h) Quantification of IGFBP2 levels from (g) (n=3; *, p < 0.05; **, p < 0.01; ns, not significant). (i) IGF1 levels measured by ELISA in astrocyte-conditioned media (ACM) from WT, KO, and KO + peptide conditions (n=4; *, p < 0.05; ***, p < 0.001; two-tailed t-test for all comparisons).

To identify additional protein complexes interacting with IGF1(1-3), we next performed a biotinylated IGF1(1-3) peptide pull-down followed by mass spectrometry (Fig. 4d). In this assay, the biotinylated peptide serves as a bait to isolate interacting and proximal proteins, enabling identification of peptide-associated protein complexes rather than strictly direct binding partners. Thus, differences in enrichment reflect changes in the protein interaction environment associated with the peptide in WT versus KO astrocytes. In WT astrocytes, peptide-associated complexes showed selective enrichment for a subset of mitochondrial-associated proteins, with the strongest signal arising from components of the dehydrogenase complex. While additional mitochondrial pathways showed a trend toward positive enrichment, these did not reach statistical significance (q > 0.05). In contrast, in KO astrocytes, proteasomal components were strongly enriched (Fig. 4e, f; SI Appendix, Dataset S7). The shift toward proteasome-associated complexes indicated that the peptide is preferentially associated with protein degradation machinery in KO astrocytes, suggesting that IGF1(1-3) peptide may promote proteasome-dependent turnover of IGFBP2.

Consistent with this hypothesis, we found that peptide treatment reduced IGFBP2 levels in KO astrocytes, and this reduction was blocked by the addition of proteasome inhibitor MG-132, confirming that IGFBP2 degradation requires proteasomal activity (Fig. 4g, h). Thus, IGF1(1-3) peptide likely acts through both mechanisms, displacing IGFBP2 from IGF1 to reduce sequestration and targeting IGFBP2 for proteasome-dependent degradation.

Finally, we tested the functional consequence of these mechanisms by measuring IGF1 levels in ACM by ELISA. KO ACM contained lesser IGF1 than WT (24.3 ± 1.7 vs. 52 ± 10.2 pg/mL). Importantly, peptide treatment significantly increased IGF1 levels in KO ACM (Fig. 4i). This indicates that IGF1(1-3) peptide not only reduces IGFBP2 abundance but also functionally rescues IGF1 bioavailability in the astrocytic environment, providing a direct link between IGFBP2 degradation and improved neuronal access to IGF1.

### IGF1(1-3) peptide reverses IGFBP2-linked mitochondrial dysfunction in MeCP2 KO astrocytes

Given the differential enrichment of proteasomal components in KO astrocytes and selective mitochondrial-associated proteins in WT astrocytes (Fig. 4e,f), we next asked whether IGFBP2 dysregulation impacts mitochondrial pathways in KO astrocytes. To address this, we evaluated the KO astrocyte proteome before and after IGF1(1-3) peptide treatment (Fig. 5a). Gene set enrichment analysis (GSEA) of KO astrocytes relative to WT revealed a significant negative enrichment of mitochondrial and metabolic pathways including cellular respiration and oxidative phosphorylation (Fig. 5a), indicating reduced representation of these pathways in KO astrocytes. Peptide treatment was able to reverse the enrichment scores for most of these pathways shifting them towards WT levels, consistent with partial rescue (Fig. 5a). Consistent with this, leading-edge analysis demonstrated that 46 of the 50 proteins contributing to the negative enrichment of oxidative phosphorylation were partially restored following peptide treatment (Fig. 5a, b; SI Appendix, Datasets S2, S6). These leading-edge proteins represent the core subset driving the pathway-level deficit observed in KO astrocytes.

**Fig. 5.**
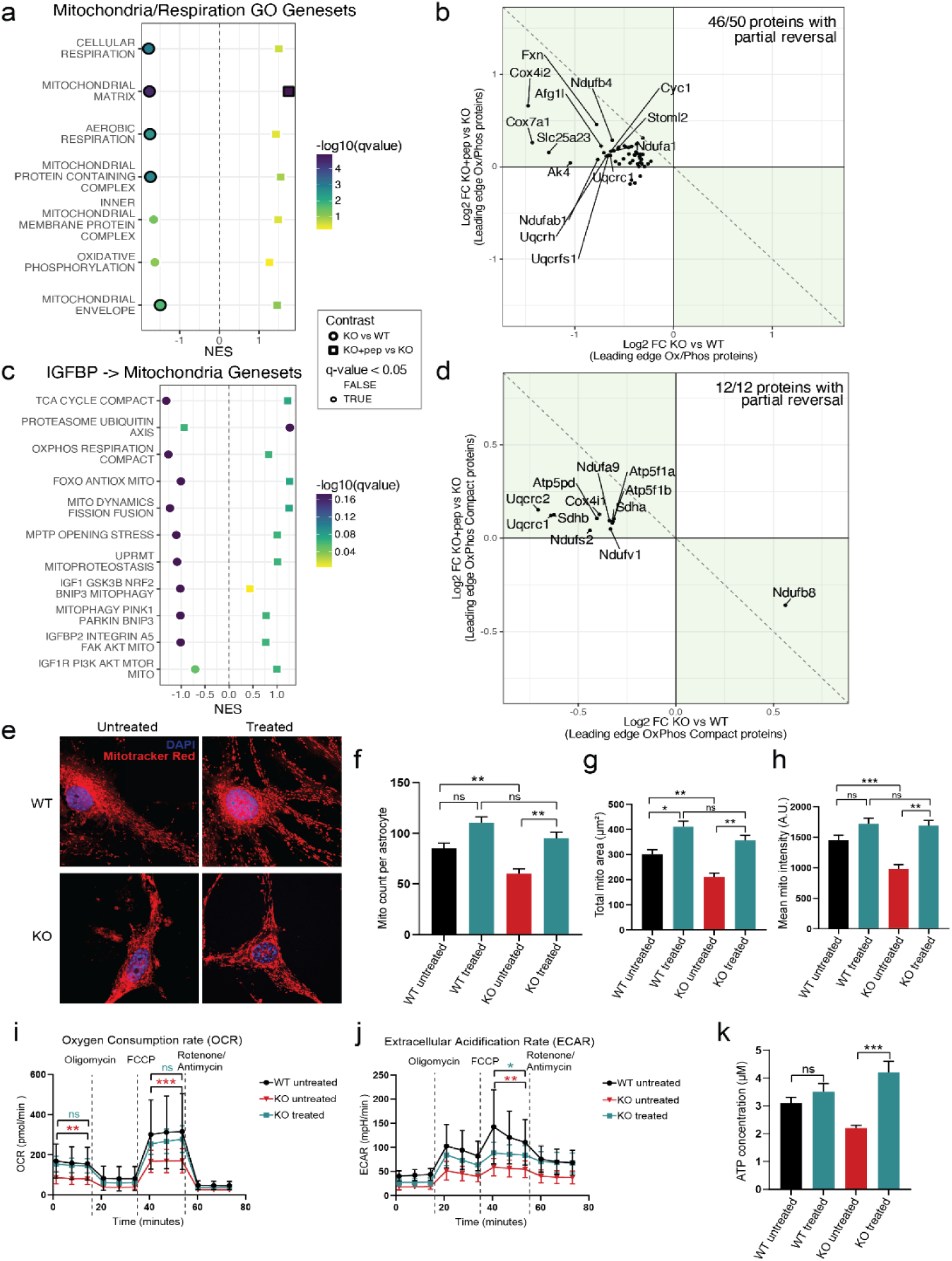
IGF1(1-3) peptide rescues mitochondrial dysfunction in MeCP2 KO astrocytes. (a) GSEA normalized enrichment score plot showing top mitochondrial-related pathways suppressed in KO vs. WT astrocytes (left) and their positive enrichment in KO astrocytes treated with peptide vs. untreated (right). Pathways include cellular respiration, mitochondrial inner membrane, mitochondrial protein-containing complex, and oxidative phosphorylation. Different shapes denote the two conditions and color indicates the q-value; x-axis represents normalized enrichment score (NES). Significant pathway enrichments are indicated with black borders. (b) Scatter plot of leading-edge oxidative phosphorylation proteins showing reversal in protein expression changes upon peptide treatment of KO astrocytes. (c) GSEA normalized enrichment score plot for IGF1 and IGFBP regulated mitochondrial pathways. Several pathways show nominal reversal of NES in KO + peptide vs. KO. (d) Leading edge analysis identifying 12/12 “oxphos compact” proteins reversed by peptide treatment. (e) Representative images of WT and KO astrocytes, untreated or peptide-treated, stained with Mitotracker Red CMXRos (mitochondria, red) and DAPI (nuclei, blue) Scale bar = 10 μm. (f-h) Quantification of mitochondrial number per cell (f), total mitochondrial area per cell (g), and mean mitochondrial fluorescence intensity (h) from Mitotracker images (n = 5 independent cultures; *, p < 0.05; **, p < 0.01; ***, p < 0.001; one-way ANOVA with Tukey’s post hoc test). (i, j) Seahorse extracellular flux analysis of oxygen consumption rate (OCR; pmol/min) (i) and extracellular acidification rate (ECAR; mpH/min) (j) in WT and KO astrocytes with or without peptide treatment. OCR and ECAR are significantly reduced in KO astrocytes and are restored toward WT levels after treatment (*, p < 0.05; **, p < 0.01; ***, p < 0.001; ns, non-significant). (k) ATP concentration in astrocyte-conditioned media (μM), showing decreased release from KO astrocytes and significant rescue upon peptide treatment (***, p < 0.001; one-way ANOVA with Tukey’s post hoc test).

Since IGFBP2 and other IGFBPs are known to affect mitochondrial structure and function, we curated several IGF1-mitochondria composite gene sets to specifically examine mitochondrial proteins that are regulated by IGF1 and IGFBP2 based on published literature (33–35). We evaluated known pathway proteins including: (i) IGFBP2-integrin/FAK-Akt signaling, (ii) PTEN-autophagy/mitophagy components, and (iii) IGF1-dependent mitochondrial protective pathways (NRF2-BNIP3, PGC-1α-NRF1/TFAM modules). These pathways showed consistent trends toward dysregulation in KO astrocytes and partial normalization following peptide treatment (Fig. 5c). In particular, leading edge analysis of the oxidative phosphorylation compact gene set revealed that 12/12 proteins were partially reversed by IGF1(1-3) treatment (Fig. 5d). These included proteins such as Ndufa9, Atp5f1a, and Atp5f1b. Notably, several of these proteins are known downstream targets of Akt and AMPK signaling (36), providing a potential mechanistic link between IGFBP2-mediated signaling and mitochondrial dysfunction in KO astrocytes.

To directly relate these proteomic shifts to mitochondrial physiology, we examined functional and morphological changes in mitochondria. MitoTracker imaging revealed reduced mitochondrial number and size, as well as mean MitoTracker intensity in KO astrocytes (Fig. 5e-h). All of these deficits were rescued in astrocytes treated with IGF1(1-3) peptide (Fig. 5e-h). Furthermore, seahorse extracellular flux analysis showed significantly decreased basal and maximal oxygen consumption rates (OCR) and extracellular acidification rate (ECAR) in KO astrocytes compared to WT. Again, peptide treatment of astrocytes restored ECAR and OCR rates to near WT levels (Fig. 5i, j). Finally, ATP bioluminescent assay showed that peptide treatment of KO-A increased extracellular ATP two-fold, rescuing ATP release to WT levels after treatment (Fig. 5k).

Taken together, these findings demonstrate that elevated IGFBP2 in MeCP2-deficient astrocytes is associated with a coordinated suppression of mitochondrial and metabolic pathways, resulting in reduced oxidative phosphorylation (OXPHOS) protein abundance, impaired mitochondrial morphology, decreased respiration, and diminished ATP release. IGF1(1-3) peptide treatment promotes proteasome-dependent degradation of IGFBP2, restores IGF1 bioavailability, and rescues mitochondrial proteomic networks and cellular bioenergetics in astrocytes.

### IGF1(1-3) peptide treatment targeted to astrocytes rescues neuronal growth and synaptogenesis by normalizing IGF1 signaling and synaptic protein expression in neurons

To determine the molecular consequences of astrocyte-targeted IGF1(1-3) peptide treatment on neurons, we performed quantitative proteomic analysis of neurons co-cultured with either untreated or peptide-treated KO astrocytes (Fig. 6a). Several proteins were differentially expressed in KO neurons co-cultured with untreated KO astrocytes compared to WT co-cultures (such as ATP receptor P2rx7), while treatment of KO astrocytes with IGF1(1-3) reversed changes in a range of proteins in KO neurons (such as postsynaptic protein Homer3) (SI Appendix, Fig. S4a, b; Dataset S8). Notably, IGF1(1-3) had no significant effect on the proteome of KO neurons cultured in isolation, confirming that the rescue is mediated importantly via astrocytes (SI Appendix, Fig. S4c).

**Fig. 6.**
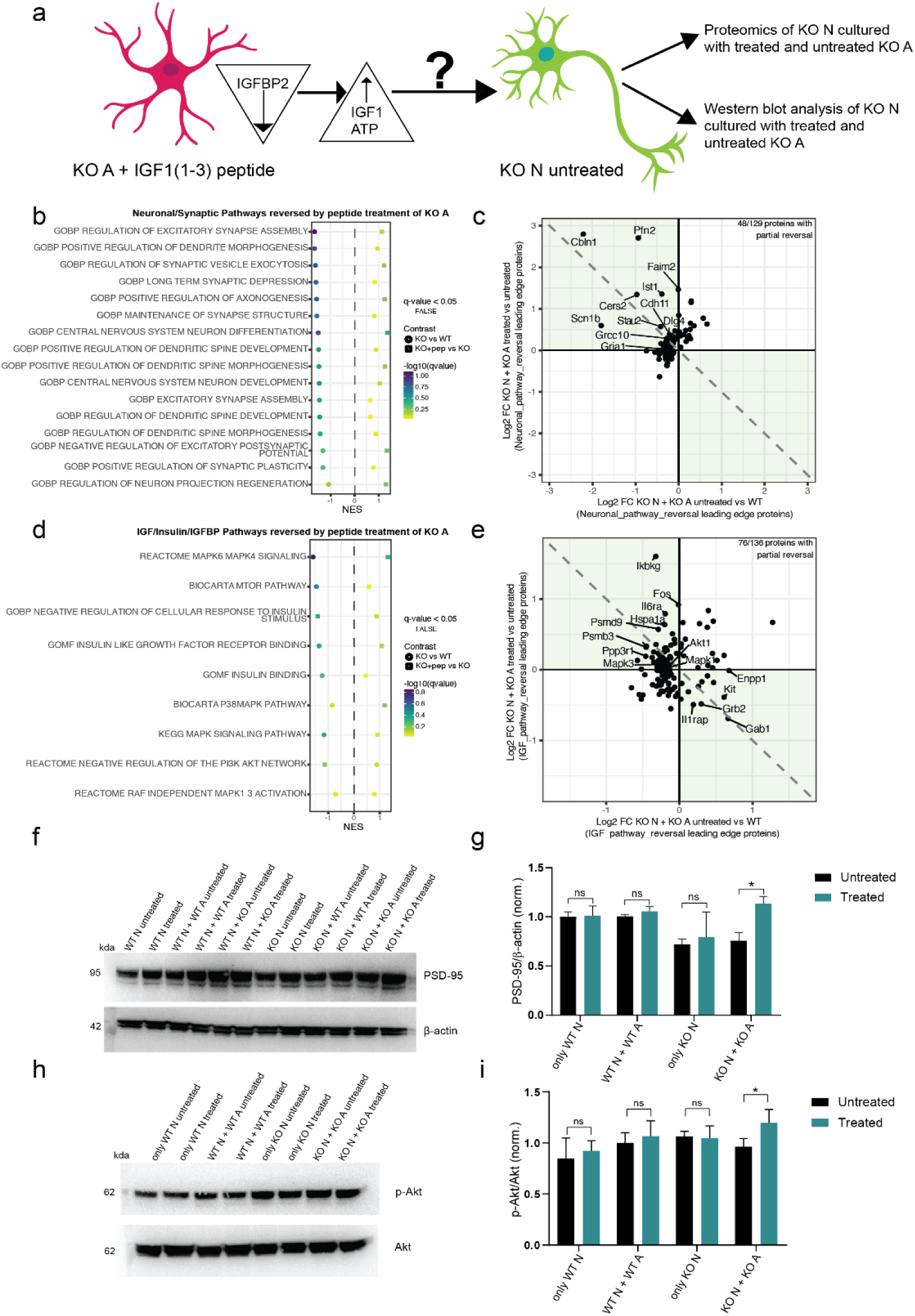
IGF1 peptide treatment of astrocytes restores synaptic protein levels and IGF1 downstream signaling pathways in neurons. (a) Schematic illustrating experimental design and logic. IGF1(1-3) treatment of KO-A leads to downregulation of IGFBP2 and increase in IGF1 and ATP levels in the media. To answer how these changes affect KO-N, they were co-cultured with treated or untreated WT- or KO-A and neuronal proteomes were analyzed to assess astrocyte-mediated effects. (b) GSEA normalized enrichment score plot of neuronal/synaptic pathways nominally reversed by peptide treatment of KO-A in co-culture, highlighting partial restoration of dendritic spine development, excitatory synapse assembly, and synaptic plasticity. (c) Scatter plot of leading-edge proteins from neuronal/synaptic pathways that exhibit partial or complete reversal following peptide treatment (48/129 proteins show partial reversal; green-shaded region). (d) GSEA normalized enrichment score plot of IGF/insulin/IGFBP-related pathways nominally reversed in neurons co-cultured with peptide-treated KO-A. (e) Scatter plot of leading-edge proteins from IGF/insulin/IGFBP-related pathways with reversal following peptide treatment (76/136 proteins show partial reversal; green-shaded region). (f-i) Western blot validation and quantification of selected neuronal proteins. (f, g) Postsynaptic density protein PSD-95 expression levels are increased in KO-N co-cultured with treated KO-A. (g, h) Phospho-Akt and total Akt levels are increased in KO-N co-cultured with treated KO-A, showing reactivation of downstream IGF1 signaling in neurons. Data represent mean ± SEM (n = 3 independent cultures; * p < 0.05, ** p < 0.01; one-way ANOVA with Tukey’s post-hoc test).

GSEA analysis comparing neurons co-cultured with peptide-treated versus untreated KO astrocytes revealed top pathways showing significant reversal based on their absolute NES scores (SI Appendix, Dataset S9). We focused on pathways associated with synapse assembly, dendritic spine formation, and IGF1-dependent signaling. While several pathways did not reach statistical significance, they still exhibited opposite enrichment trends (Fig. 6b, d). Neurons co-cultured with untreated KO astrocytes displayed suppression of pathways essential for excitatory synapse assembly, dendritic spine morphogenesis, and synaptic plasticity (Fig. 6b). Treatment with IGF1(1-3) showed activation of these pathways, with key reversed leading-edge proteins including DLG4 (PSD-95) and GRIA1 (AMPA receptor subunit) (Fig. 6c).

Further, GSEA of these expression changes also demonstrated reversal in enrichment scores in pathways related to IGF1/IGF1R signaling, with leading-edge proteins such as AKT1, MAPK3, and MAPK1 (ERK1/2) normalized following peptide treatment (Fig. 6d, e). These changes are particularly relevant, as Akt and ERK1/2 are critical downstream effectors of the IGF1/insulin pathway. Additional proteins showing reversed fold changes included IGF1R (IGF1 receptor), AKT2, NTRK2 (BDNF receptor), and DLGAP1 (postsynaptic density protein) (SI Appendix, Fig. S4d). Western blot validation confirmed increased PSD-95 and phosphorylated Akt (p-Akt) levels in neurons co-cultured with treated KO astrocytes, consistent with restoration of postsynaptic integrity and downstream IGF1 pathway activation (Fig. 6f-i).

These data demonstrate that IGF1(1-3) peptide treatment, by targeting MeCP2-deficient astrocytes, broadly and at least partially normalizes the neuronal proteome, restoring fundamental developmental programs, rebalancing key signaling pathways such as PTEN/PI3K/Akt, and rescuing synaptic and mitochondrial protein networks.

Finally, to test the therapeutic relevance of these findings *in vivo*, we administered IGF1(1-3) peptide daily (0.02 mg/g, i.p.) from postnatal day 18-28 in WT, global MeCP2-null (MeCP2^-/y^), and astrocyte-specific conditional KO mice (Aldh1l1-Cre; MeCP2^-/y^) (SI Appendix, Fig. S5a). Open field tests showed that locomotor activity deficits in both models were significantly improved with peptide treatment (SI Appendix, Fig. S5b, c). In rotarod assays, motor learning deficits were partially rescued in astrocyte-specific KO mice but remained impaired in global nulls (SI Appendix, Fig. S5d), suggesting that some motor deficits can be ameliorated by IGF1(1-3) peptide potentially targeting astrocytic MeCP2, whereas others may be refractory to such treatment.

Taken together, our findings support a model in which MeCP2-deficient astrocytes upregulate IGFBP2, which sequesters IGF1 and limits neuronal PI3K-Akt signaling, PSD95 expression, and excitatory synaptogenesis. Treatment with the IGF1(1-3) peptide hampers IGFBP2 interaction with full-length IGF1 and promotes its proteasomal degradation, thereby restoring IGF1 bioavailability to neurons. Importantly, our proteomics and Western blot analyses indicate that this dual mechanism restores both neuronal signaling (PI3K/Akt/mTOR, PSD95 levels) and astrocytic metabolic support (mitochondrial structure, OXPHOS proteins, ATP release) (Fig. 7).

**Fig. 7.**
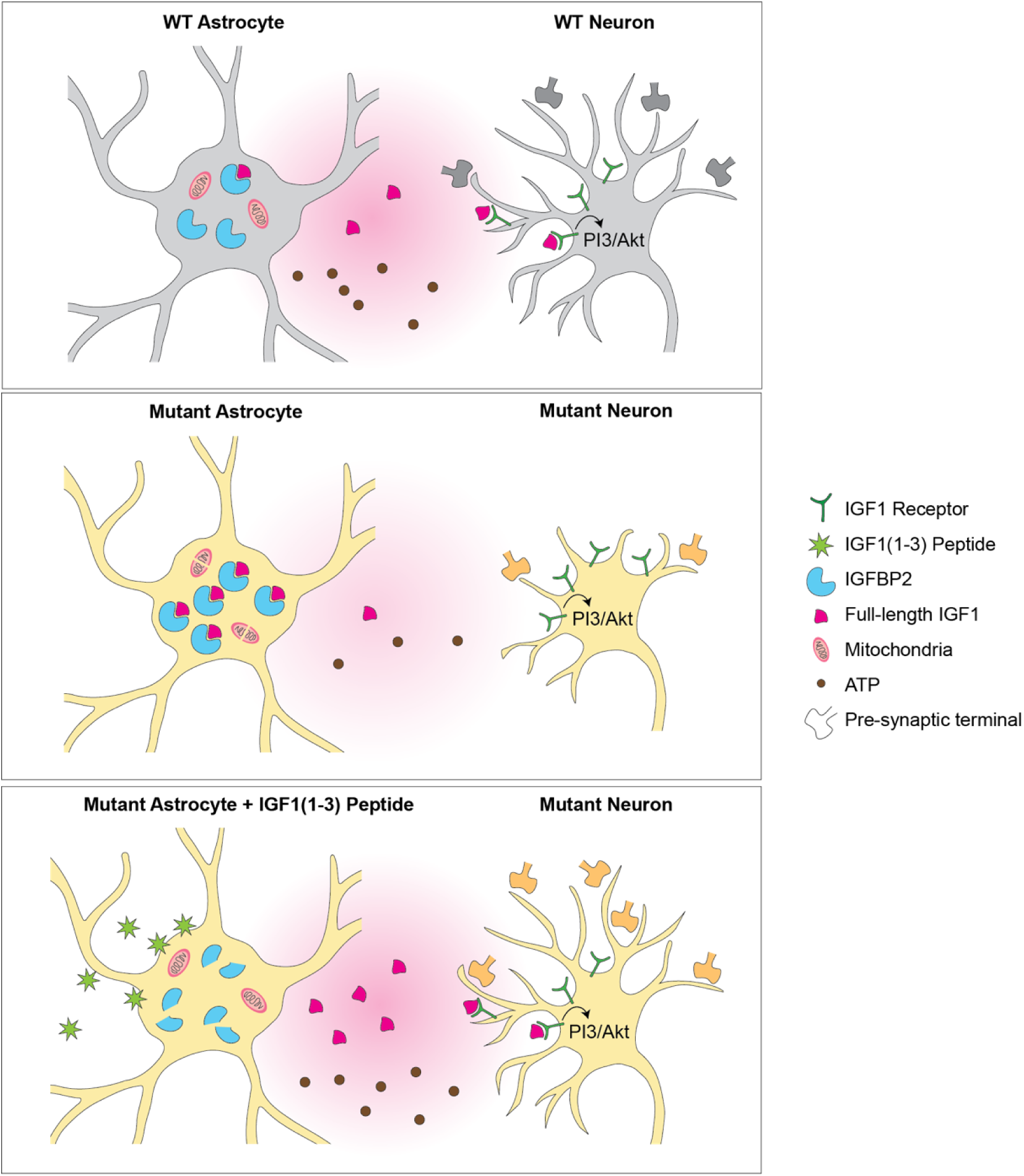
Proposed model of astrocyte-mediated synaptic and mitochondrial dysfunction in MeCP2 deficiency and rescue by IGF1(1-3) peptide. WT astrocytes secrete IGF1 and ATP that supports neurite outgrowth and synaptogenesis in WT neurons. KO astrocytes overexpress IGFBP2, that sequesters IGF1 and breaks down mitochondria, thus impairing neuronal PI3K-Akt signaling and synaptogenesis. Loss of mitochondrial integrity and function in KO astrocytes is accompanied by decreased ATP production. IGF1(1-3) peptide treatment of KO astrocytes degrades IGFBP2 and restores mitochondrial integrity and function, thus rescuing ATP release and IGF1 bioavailability, which in turn rescues neuronal PI3/Akt signaling and restores synapse formation.

## Discussion

Rett syndrome (RTT) is a devastating neurodevelopmental disorder caused by loss-of-function mutations in *MECP2* (37–40). While traditionally considered a neuron-centric disorder, growing evidence highlights astrocytes as critical non-cell-autonomous regulators of RTT pathology (24, 25, 29, 41). Here, we identify astrocyte-secreted IGFBP2 as a molecular brake on synaptogenesis regulated by MeCP2-deficient astrocytes and show that treatment with IGF1(1-3) peptide degrades IGFBP2, restores IGF1 availability, rescues mitochondrial and metabolic support, and normalizes neuronal PI3K-Akt-PSD95 signaling. These findings provide a mechanistic explanation for the efficacy of IGF1(1-3) and potentially for its mimetic Trofinetide, the first FDA-approved drug for RTT (20).

The bidirectional regulation of IGFBP2 and IGF1 signaling illustrates the dual roles of MeCP2 as transcriptional repressor and activator (5–7, 9). In astrocytes, MeCP2 mutations cause increased expression of IGFBP2 (29; this study), consistent with the role of MeCP2 as transcriptional repressor. At the same time, IGF1 signaling and its downstream pathway in neurons is downregulated, as is BDNF signaling (13, 42–44), consistent with the role of MeCP2 as transcriptional activator. The reduction of IGF1 signaling is at least partially due to increased IGFBP2, which binds IGF1 and reduces its availability, and supplying exogenous IGF1 can partially rescue symptoms and IGF1 signaling in neurons of Rett model mice (14, 30). While the application of IGF1 peptide also partially rescues symptoms and IGF1 signaling (13), an open question has been why IGF1(1-3) peptide mimics the effects of IGF1 despite evidence that it does not engage the IGF1 receptor (IGF1R) with the same affinity or in the same manner as full-length IGF1 (45, 46). Our study provides a plausible explanation: rather than acting directly on neurons, IGF1(1-3) promotes proteasomal degradation of IGFBP2 in astrocytes. By removing this inhibitory binding protein, IGF1 is made available to engage neuronal receptors and trigger canonical PI3K-Akt-S6 and PSD95/GluA1 signaling cascades. This mechanism clarifies how IGF1(1-3) converges functionally with IGF1 on synaptic outcomes, despite differences in receptor interactions.

Our study also contributes to a growing body of work exploring the mechanisms by which astrocytes regulate neuronal development. Consistent with previous studies showing that astrocytes secrete synaptogenic molecules such as thrombospondins, hevin, and glypicans (47, 48), we find that MeCP2-deficient astrocytes also secrete synapse-suppressive cues, including IGFBP2 and other ECM regulators. It is reasonable to propose that IGFBP2 upregulation sequesters IGF1, disrupts PI3K-Akt signaling, and leads to a loss of PSD-95 and other synaptic proteins in neurons. We show that IGF1 peptide treatment rescues PSD-95 levels, Homer3 expression, and Akt signaling, suggesting that the peptide restores not only synaptic structure but also functional IGF1 signaling cascades. These findings align with *in vivo* observations that IGF1 analogs improve synaptic function, plasticity and behavioral outcomes in RTT mouse models (13, 14) and provide direct mechanistic evidence of how astrocyte-mediated IGF1 signaling modulates neuronal circuit assembly.

Interestingly, our study did not find significant changes in inhibitory synapse formation as shown by gephyrin/Bassoon colocalization across genotypes or treatment conditions (SI Appendix, Fig. S1e, f; Fig. S1l, m). Similar effect was seen previously upon IGF1(1-3) treatment of neurons (49). This suggests that astrocytic MeCP2 loss preferentially affects excitatory synapse development, possibly due to selective regulation of excitatory synaptogenic proteins. These results support the excitation-inhibition imbalance seen in RTT (50), based on reduction of excitatory synaptic transmission to both excitatory and inhibitory neurons (51). It is also possible that inhibitory circuits may be less sensitive to IGFBP2-mediated IGF1 sequestration, or the time frame of treatment (24 h) may not be sufficient to induce observable changes. These observations warrant deeper investigation into the circuit- and cell-type-specific responses to astrocyte-mediated IGF1 signaling.

Our data also highlight a second dimension of astrocytic dysfunction, viz. impaired mitochondrial support. MeCP2-deficient astrocytes displayed fragmented mitochondria, reduced oxygen consumption, and diminished ATP release. These physiological deficits map directly onto suppressed expression of mitochondrial OXPHOS proteins and regulators (e.g., Ndufs6, Atp5a1, Cox6b1, Mfn2, Mdh1, Idh1), which were consistently downregulated in KO astrocytes and restored by peptide treatment. Many of these proteins are direct Akt or AMPK targets, positioning IGFBP2-integrin/FAK-Akt signaling as a mechanistic link between IGFBP2 dysregulation and mitochondrial health (30, 33, 36, 52). While our functional assays focused on OCR and ATP release, the concordant reversal of OXPHOS proteins by IGF1(1-3) provides molecular evidence that mitochondrial function is directly tied to IGFBP2 regulation. This integrated path from proteomics to physiology strengthens the case that astrocytic metabolism is an essential contributor to RTT pathology.

Interestingly, mitochondrial proteins were among the top interactors of IGF1(1-3) in WT astrocytes but were replaced by proteasomal proteins in MeCP2 KO astrocytes (Fig. 4e, f), suggesting a shift in cellular priorities from energy production to proteostasis. Previous studies have implicated mitochondrial dysfunction in RTT pathology (27, 53, 54), and our work expands this view by positioning astrocytes, and specifically IGFBP2 dysregulation, as a contributing factor. Notably, neutralizing IGFBP2 in RTT ACM has been shown to rescue dendritic outgrowth in neurons (29), supporting our conclusion that astrocyte-secreted IGFBP2 is a core molecular brake on neuronal maturation. Protein accumulation is a hallmark of several neurodevelopmental and neurodegenerative disorders, and our study suggests that IGF1(1-3) peptide and Trofinetide may also recruit the proteasomal machinery and restore protein balance.

Notably, the astrocyte-specific conditional knockout experiments support the concept of astrocytes as therapeutic targets for neurodevelopmental and other disorders. IGF1(1-3) peptide treatment improved locomotion and partially rescued motor learning in these mice, whereas global MeCP2-null mice showed only partial behavioral rescue. Indeed, IGF1(1-3) treatment has been shown previously to improve a range of phenotypes including breathing variability, heart rate, locomotion and life span in MeCP2-null mice (13). Prior work showed that genetic restoration of MeCP2 in astrocytes improved locomotion and breathing patterns in MeCP2-null mice (26). Thus, several key phenotypes can be ameliorated through astrocytic rescue, though different brain regions that mediate specific functions may be differentially amenable to such rescue.

Our study has several limitations. First, although our proteomics and functional assays established a strong link between IGFBP2 and mitochondrial support, more detailed biochemical validation (e.g., Western blots or immunostaining for individual OXPHOS proteins) would strengthen the pathway-level claims. Second, we did not directly quantify astrocyte- versus neuron-specific IGFBP2 levels *in vivo* before and after peptide treatment. Third, while our *in vivo* experiments provide proof-of-concept behavioral rescue, they were limited in scope; future work should expand to systemic and behavioral endpoints which are clinically relevant in RTT.

Together, these findings identify astrocytic IGFBP2 as a central mediator of non-cell-autonomous dysfunction in Rett syndrome and provide a mechanistic explanation for the efficacy of IGF1(1-3) peptide in treating RTT. By promoting proteasomal degradation of IGFBP2, the peptide enhances IGF1 signaling to neurons and normalizes astrocytic mitochondrial support, thereby rescuing both synaptic and metabolic deficits. This dual action highlights astrocytes not simply as passive contributors but as active gatekeepers of neuronal circuit maturation. From a translational perspective, IGFBP2 is potentially both a therapeutic target and a biomarker of treatment response. More broadly, our work emphasizes that targeting astrocytic dysfunction is sufficient to restore neuronal signaling and behavior, supporting the paradigm of cell-type-specific interventions for neurodevelopmental disorders. Future studies aimed at astrocyte-selective delivery of IGF1 mimetics, or direct modulation of IGFBP2 and its downstream signaling pathways, may thus open new avenues for durable, mechanism-based therapies for Rett syndrome and related conditions.

## Materials and Methods

### Animals

All animal experiments were approved by the Institutional Animal Care and Use Committee (CAC) at the Massachusetts Institute of Technology and conducted in accordance with the National Institutes of Health guidelines. Wild-type (WT) and Mecp2-null mice (B6.129P2(C)-Mecp2^tm1.1Bird/J^, Stock #: 003890) were obtained from The Jackson Laboratory. Astrocyte-specific conditional knockout (cKO) mice were generated by crossing floxed Mecp2 mice with Aldh1l1-Cre mice (The Jackson Laboratory, Stock #: 023748). Both male and female pups were used for *in vitro* experiments; only male animals were used for *in vivo* behavioral studies.

### Primary Cell Culture

*Astrocyte Cultures.* Cortical astrocytes were isolated from postnatal day 1-3 (P1-P3) WT or Mecp2-null mice. Cortices were dissected, meninges removed, and tissue dissociated in 0.05% trypsin-EDTA (Thermo Fisher). Cells were plated in Astrocyte media (ScienCell Technologies) with 10% fetal bovine serum (FBS) and 1% penicillin-streptomycin and maintained for 10-14 days. Confluent monolayers were shaken at 130-150 rpm in an orbital shaker to remove microglia and oligodendrocyte precursor cells, then replated for downstream assays. Pups were genotyped by PCR prior to tissue collection.

*Neuron-Astrocyte Co-culture System.* Cortical neurons were isolated from P0 mice and seeded on poly-D-lysine-coated coverslips at a density of 125,000 cells per well in 24-well plates. After 3 d *in vitro* (DIV3), fluorodeoxyuridine (FuDr) (Sigma) was added to inhibit glial proliferation. For indirect co-culture, astrocytes were plated on transwell inserts in Neurobasal medium (Thermo Fisher) and suspended above neuronal monolayers. Co-culture was maintained for 7-10 d in Neurobasal medium supplemented with B27, GlutaMAX, and antibiotic-antimycotic. For all experiments, four conditions were used: WT neurons (WT-N) + WT astrocytes (WT-A), WT-N + KO-A, KO-N + WT-A, and KO-N + KO-A (Fig. 1a). All the combinations were analyzed, representative conditions are shown for clarity and to avoid overcrowding of the figure. IGF1(1-3) peptide treatment was performed at DIV7 for 24 h prior to setting up the co-cultures.

### Peptide Treatment

A synthetic IGF1(1-3) peptide (Gly-Pro-Glu; GPE) was obtained from Bachem and reconstituted in sterile water. Astrocytes or neurons were treated with 100 ng/mL peptide for 24 h unless otherwise specified. Co-cultures were established 24 h after peptide treatment. To control for potential effects of IGF1(1-3) peptide diffusion across the transwell membrane, only WT- or KO-treated astrocyte or neuron (A or N) conditions were used. Dose-response experiments (50-200 ng/mL; SI Appendix, Fig. S1i, j) confirmed that 100 ng/mL was appropriate for all functional assays. For proteasome inhibition, cells were co-treated with MG-132 (10 µM; Sigma-Aldrich) for 4 h.

### Immunocytochemistry and Spine Analysis

Cells were fixed with 4% (vol/vol) paraformaldehyde (PFA), permeabilized with 0.1% Triton X-100, and blocked with 5% (vol/vol) normal goat serum. Primary antibodies included rabbit anti-MAP2 (1:500; Abcam), mouse anti-GFAP (1:500; Millipore), rabbit anti-PSD-95 (1:500; Sigma), guinea pig anti-Bassoon (1:500; Synaptic Systems), mouse anti-Gephyrin (1:500; Synaptic Systems), and rabbit anti-GFP (1:500; Abcam). Species-specific secondary antibodies conjugated to Alexa Fluor dyes (Thermo Fisher) were used. Images were acquired on a Leica SP8 confocal microscope and analyzed in FIJI/ImageJ. Spine number and density were quantified along secondary dendrites over 50-μm segments from 10-15 neurons per condition (Fig. 1c, d).

### Western Blotting

Cells or tissues were lysed in RIPA buffer containing protease and phosphatase inhibitors. Protein concentration was determined by BCA assay. Equal protein amounts were separated by SDS/PAGE, transferred to PVDF membranes, and probed with antibodies against MeCP2 (1:1000; Cell Signaling Technology), PSD-95 (1:1000; Cell Signaling Technology), IGFBP2 and other IGFBPs (1:1,000; R&D Systems), Phospho-p44/42 MAPK (Erk 1/2) (1:1000; Cell Signaling Technology), FLAG (1:200; Sigma), p44/42 MAPK (Erk 1/2) (1:1000; Cell Signaling Technology), Phospho-Akt (Ser473) (1:1000; Cell Signaling Technology), Akt (1:1000; Cell Signaling Technology), and β-actin (1:2000; Sigma). Blots were visualized by ECL (Millipore) on a ChemiDoc (Bio-Rad) and quantified by densitometry in ImageJ.

### Coimmunoprecipitation (Co-IP)

pLV CMV IGF1-FLAG was a gift from Larry Gerace (Addgene plasmid # 175170). WT and KO astrocytes were transfected with the full-length FLAG-tagged IGF1 expression plasmid using Lipofectamine 3000 (Thermo Fisher). After 24 h of treatment with 100 ng/mL IGF1(1-3) peptide, cells were lysed, and Co-IP was performed using anti-FLAG M2 agarose beads (Sigma). Precipitates were washed, eluted, and probed for IGFBP2 by Western blot.

### Proteomics

#### Sample Preparation

Conditioned media (ACM) and astrocyte lysates were collected after 24 h of serum-free incubation (with or without peptide treatment). Samples were concentrated and protein was digested using trypsin. Peptides were desalted using C18 spin columns and submitted for DIA or TMT analysis.

#### Data-Independent Acquisition (DIA) Mass Spectrometry

Samples were analyzed on an Orbitrap Eclipse mass spectrometer coupled to a Vanquish Neo UHPLC system (Thermo Fisher). Peptides were separated on a 45-min gradient. MS data were acquired in DIA mode. Data were processed using DIA-NN (v1.8) in library-free mode against the Mus musculus UniProt database.

#### Tandem Mass Tag (TMT) Mass Spectrometry

For TMTpro experiments, peptides were labeled, fractionated, and analyzed on an Orbitrap Eclipse. Data were processed using PEAKS Studio (v10.6).

#### Data Analysis

Protein intensity values were log2-transformed and normalized. Differential expression analysis was performed using limma. Gene set enrichment analysis (GSEA) overrepresentation analyses were performed using clusterProfiler (v.4.8.2) with pathway sets from Gene Ontology (GO), KEGG, and Reactome, downloaded through msigdbr (v.7.5.1).

Detailed description of methods is provided in SI Appendix.

### Biotinylated Peptide Pull-Down and Mass Spectrometry

Biotinylated IGF1(1-3) peptide was synthesized (Genscript) and incubated with Dynabeads streptavidin magnetic beads (ThermoFisher). Astrocyte lysates were incubated with bead-bound peptide, and bound proteins were digested on-bead. Peptides were labeled with TMTpro, fractionated, and analyzed by LC-MS/MS. Data were processed as described for TMT proteomics. Detailed description of methods is provided in SI Appendix.

### Mitochondrial Imaging and Functional Assays

#### Mitotracker Staining

Cells were stained with MitoTracker Red CMXRos (100 nM, Thermo Fisher) and DAPI (1 μg/mL) for 30 min at 37°C. Images were acquired by confocal microscopy. Mitochondrial area, intensity, and count were quantified using custom ImageJ macros.

#### Seahorse Metabolic Assay

Oxygen consumption rate (OCR) and extracellular acidification rate (ECAR) were measured using a Seahorse XF Analyzer (Agilent) following the Mito Stress Test protocol. Parameters included basal respiration, ATP production, and spare respiratory capacity.

#### ATP Quantification

Extracellular ATP levels were measured using a luciferase-based assay (ATP Bioluminescence Kit, Abcam) and normalized to cell number.

### ELISA

IGF1 levels in ACM were measured using a mouse IGF1 ELISA kit (Thermo Fisher) according to the manufacturer’s instructions. Samples were run in technical duplicates and analyzed on a plate reader at 450 nm.

### RNA Isolation and qPCR

RNA was extracted (RNeasy kit; Qiagen), and cDNA was synthesized (Takara). qPCR was performed using SYBR Green Master Mix on a QuantStudio system. Gene expression was normalized to *Gapdh* (ΔΔCt method).

### Behavioral Studies

Male Mecp2-null and Aldh1l1-Cre;Mecp2^flox/y^ mice received daily i.p. injections of IGF1(1-3) peptide (0.02 mg/g) or vehicle from P18 to P28. Behavioral tests were conducted at P29-30.

*Open Field Test.* Locomotor activity was tracked for 15 min using Bonsai software.

*Rotarod.* Mice were tested on an accelerating rotarod (4-40 rpm over 5 min) over 3 days; latency to fall was recorded.

### Statistical Analysis

Data are presented as mean ± SEM. Group comparisons were made using unpaired two-tailed Student’s t-tests or one-way ANOVA followed by Tukey’s post hoc test. p < 0.05 was considered statistically significant. Statistical analyses were performed using GraphPad Prism.

## Acknowledgements

We thank Taylor Johns for technical and administrative support. We thank Jiho Park for assistance in preparing Fig. 7. We thank Brooke Linnehan from the Whitehead proteomics core for her assistance with the proteomics experiments. This research was supported by NIH training grant T32MH112510 (R.M.L.), International Rett Syndrome Foundation (D.L.T), NIH grant RF1AG075901 (E.F.), NIH grant 5R01MH104610 (R.J.), NIH grant R01MH085802 and the Simons Foundation Autism Research Initiative (SFARI) (M.S.).

## Author Contributions

Conceptualization: P.O., R.J., M.S.; Methodology: P.O., V.K., F.S.; Investigation: P.O., A.B., F.S., R.M.L.; Formal analysis: P.O., V.K., F.S.; Visualization: P.O., V.K.; Resources: V.K., D.L.T., M.S.; Funding acquisition: R.J., M.S.; Supervision: E.F., R.J., M.S.; Writing-original draft: P.O., V.K.; Writing-review & editing: P.O., V.K., R.M.L., R.J., M.S.

## Data Sharing Plan

All data, documentation, and code used in analysis will be made available upon reasonable request.

